# Soil phosphate availability modulates the arbuscular mycorrhizal fungal community and mycorrhizal nutrition in wheat

**DOI:** 10.1101/2025.08.08.669323

**Authors:** Margot Trinquier, Eric Lecloux, Patrick Bruno, Virginie Gasciolli, Claire Jouany, Christophe Roux, Benoit Lefebvre, Agnès Ardanuy

## Abstract

- Understanding the contributions of arbuscular mycorrhizal fungi (AMF) to plant nutrition is essential for sustainable agriculture. We hypothesised that soil phosphorus (P) availability modulates the diversity and functionality of wheat root-associated AMF community, particularly the mycorrhizal nutrition.
- Wheat plants were sampled over two campaigns (2019 and 2022) in a long-term P fertilisation trial. The expression of the wheat transporters involved in the mycorrhizal nutrition was assessed by RT-qPCR, and AMF community composition by metabarcoding. Complementary experiments under controlled conditions were performed to further study the interaction between nitrogen (N) and P availability on AM.
- Different effects of P on AMF colonisation and transporter expression were observed in field-grown wheat depending on the campaigns, which differed in wheat N status. Controlled experiments confirmed that AMF colonisation depends on the limitation of either P or N, but that regulation of peri-arbuscular phosphate, ammonium and nitrate transporters depends on the nature of the limiting soil nutrients. Additionally, AMF communities varied according to the soil P availability, with the *Funneliformis* genus becoming more dominant under high P conditions for both years.
- Together, our findings show that N and P availability jointly shape root AMF communities and AM functioning. Combining community profiling and molecular markers of colonisation and nutrition offers a framework to better understand AMF contributions to plant nutrition across agroecosystems.

## Introduction

Nutrient availability in soil is a critical factor in plant productivity, influencing not only plant physiology but also plant interactions with beneficial soil microorganisms, such as arbuscular mycorrhizal fungi (AMF)(Smith & Smith, 2011). AMF associate with roots of the majority of land plants, including most crops, and provide a variety of services to host plants in exchange for carbon, such as the delivery of nutrients acquired through their vast hyphal networks, stress tolerance and pathogen resistance (Brundrett, 2002; Pozo & Azcón-Aguilar, 2007; Bennett & Groten, 2022).

This symbiosis has been shown to enhance plant nutrient acquisition under conditions of nutrient limitation, with AMF playing a crucial role in the uptake and delivery of phosphorus (P), nitrogen (N), potassium (K), sulphur, and micronutrients, such as iron, copper, manganese, and zinc (Hodge & Fitter, 2010; Wipf *et al*., 2019). Mycorrhizal plants can acquire nutrients through two distinct pathways: (1) a direct pathway, in which nutrients are acquired through the root epidermis and root hairs, and (2) a mycorrhizal pathway, in which nutrients are acquired through the AMF extraradicular mycelium and transferred to the root cortical cells containing AMF arbuscules, typical branched structures involved in plant-AMF nutrient exchanges (Smith *et al*., 2003). The relative contribution of each pathway to plant nutrition has been shown to be dependent on various biotic and abiotic factors, including soil nutrient availability, pH, soil texture, plant genotype, and the identity of AMF taxa (Smith *et al*., 2004; Cavagnaro *et al*., 2005; Feddermann *et al*., 2010; Smith *et al*., 2011).

Soil nutrient enrichment is expected to reduce the cost-benefit balance of the symbiosis in terms of nutrients acquired by a plant via the mycorrhizal pathway *vs* the carbon allocated to the AMF (Johnson, 2010; Olsson *et al*., 2010; Bennett & Groten, 2022; Bunn *et al*., 2024). Therefore, intensive mineral fertilisation practices in agriculture can disrupt AMF colonisation and nutrient exchange and potentially lead to reduced AMF biomass and diversity in soils (Treseder & Allen, 2002; Oehl *et al*., 2004; Egerton-Warburton *et al*., 2007; Oehl *et al*., 2010; Verbruggen *et al*., 2010; Edlinger *et al*., 2022). This reduction in diversity in soils potentially compromise the functional potential of AMF communities, as less diverse fungal communities may be less able to provide a full range of ecosystem functions and consequently less benefits to host plants (Powell & Rillig, 2018). Long-term fertilisation with N, P, and K has been reported to increase AMF alpha diversity in wheat roots (Ma *et al*., 2021), while it was shown that both AMF richness and phylogenetic diversity in sorghum roots were reduced when increasing short-term P fertilisation (Frew *et al*., 2023). These findings highlight the complexity of AMF responses to fertilisation and underscore the role of both environmental and host filtering in the assembly of functionally distinct communities.

In addition to the well-known effects of fertilisation on root colonisation levels and root/soil AMF community structure and composition, it is essential to understand how fertilisation influences the plant mycorrhizal nutrition. One promising approach is to track the expression level of genes coding for arbuscular mycorrhiza (AM)-specific nutrient transporters as a way to approximate nutritional benefit in ecologically realistic conditions. These transporters located on the peri-arbuscular membrane, mediate plant nutrient uptake through the mycorrhizal pathway. (Gamper *et al*., 2010; Wipf *et al*., 2019). By characterising the expression of these genes, it is possible to compare the activation of the mycorrhizal pathway for specific nutrient acquisition at a given time across contrasting fertilisation treatments, AMF communities and environmental conditions (Feddermann *et al*., 2010; Wipf *et al*., 2019).

The aim of this study was to investigate the effect of long-term P fertilisation on AMF communities and mycorrhizal nutrition in wheat. We hypothesised that (i) increasing soil P availability would reduce AMF diversity and shift fungal community composition towards species adapted to high P environments; (ii) wheat mycorrhizal nutrition would decline under high P conditions, as the plant carbon cost of maintaining AM would exceed the benefits of nutrient acquisition. To test these hypotheses, we combined the characterisation of fungal communities and wheat mycorrhizal nutrition in a 50-year P fertilisation trial during two distinct years on wheat. Differences observed between the two field campaigns led us to hypothesise that plant N status modulates AMF functioning across P availability gradients. To explore this further, we conducted controlled experiments to examine the interaction between N and P availability on AM symbiosis. By combining AMF community profiling with analysis of AM-specific nutrient transporter expression in wheat roots, we aimed to link wheat plant investment in the mycorrhizal nutrition with shifts in root-associated AMF communities under contrasting soil nutrient availability.

## Methods and Materials

### Long-term P fertilisation field trial and root sampling

A long-standing phosphorus (P) fertilisation field trial (Colomb *et al*., 2007) is maintained since 1968 at the experimental unit “Crop Phenotyping and AgroEcology Experimental Facility” (https://doi.org/10.15454/1.5483266728434124E12) in Auzeville-Tolosane, South-Western France (43.31487; 1.30185). The trial is situated on deep alluvial silty clay soil. The experimental design follows a split-plot design consisting of six randomised blocks of P treatment plots, each measuring 6 m x 15 m, with a 1 m buffer zone to prevent P dispersion. Three P treatments were evaluated: (1) No P, with no P fertilisation since 1968; (2) Low P, where P fertilisation balances annual P export from crop harvest; and (3) High P, where P fertilisation was fourfold (1968–1993), then threefold (1994– present) that of Low P, gradually increasing soil P availability. Since 1994, the P input was supplied as Triple Superphosphate adjusted to 11 and 33 kg P ha^−1^ yr^−1^ for Low P and High P treatments, respectively. In 1968, the mean P Olsen was 6.2 mg/kg in the topsoil (0-30 cm depth) and in 2021, it has decreased to 5.3 mg/kg in the No P treatment, while it has increased to 10.0 and 48.0 mg/kg in Low P and High P treatments, respectively. Soil characteristics for each plot are described in Table S1. Durum wheat (*Triticum turgidum* L. subsp. *durum* var. RGT Voilur) was cultivated over two seasons, from November to June/July in 2018–2019 and 2021–2022, at a density of 300 seeds m^−2^. Soil preparation involved superficial tillage of the topsoil layer a month prior to sowing. Nitrogen (N) fertilisation, using 33.5% ammonium nitrate, was applied to achieve a total of 174 N units in 2019 and 160 N in 2020, through 3 applications. Note that the 2019 sampling (May 2019) was done after the third ammonium nitrate application, while the 2022 sampling (April 2022) was done before the third application. In both cases, sampling was performed before heading. The technical itineraries for each campaign, sampling dates and meteorological data are summarised in Table S2 and Table S3. Root samples were collected just before the heading stage, during the stem extension phase (5 samples per plot in 2019, and 3 per plot in 2022). Each sample consisted in a pool of at least three plant root systems to a depth of 15 cm. Root samples were hand-washed with tap water to remove adhering soil, using 180 µm sieves to collect fine root fragments and immediately frozen in liquid N and stored at -80°C for subsequent DNA and RNA extraction. In May 2022 a second sampling was conducted to evaluate AMF colonisation and collect soil for the controlled condition *experiment 2*. The roots N content (%) was analysed at the plot level (by pooling the plot samples) using an Elementar Analyser (Vario ElCube).

### Experiments in controlled conditions

Experiments were carried out in a climatic chamber at 25°C and a photoperiod of 16:8 light:dark. Durum wheat seeds were rehydrated in deionised water for an hour and were then germinated on petri dishes containing gelose in the dark at 4°C for 1 week followed by four days at 28°C.

#### Experiment 1 - Testing specificity and efficiency of primers targeting the wheat AM-marker genes

Pregerminated seedlings (var. RGT Voilur) were placed in 50 mL pierced falcon tubes filled with attapulgite (Oil Dri UK) and watered with a 0.5x modified Long Ashton nutrition solution (containing 7.5 μM NaH_2_PO_4_). Half of the tubes were inoculated (Myc+) with 200 spores of *Rhizophagus irregularis* DAOM 197198 (Agronutrition) and the other half (Myc-) were left as uninoculated controls. After 5 weeks, root systems were cleaned and immediately frozen with liquid N and stored at -80°C for subsequent RNA extraction.

#### Experiment 2 - Complementary assays to long-term P fertilisation field experiments

To test whether the N:P interaction influences mycorrhizal nutrition in wheat, a factorial experiment was conducted using field trial soil from the No P and High P treatments in combination with an N fertilisation treatment. Briefly soils obtained from all the No P and High P plots were pooled according to treatment, homogenised and placed into 350 ml pots. Sterile pre-germinated wheat seedlings (var. RGT Voilur) were transplanted into No P and High P soil pots and cultivated under two nutrient treatments: (1) control (deionised water), and (2) N addition (2000 µM KNO_3_) for 10 weeks (n=7-9 pots per treatment). At the end of the experiment, root systems were washed and split in two for AMF colonisation assessment (immediately placed in KOH 10%) and for RNA extraction (immediately frozen with liquid N and stored at -80°C).

#### Experiment 3 - Evaluation of P and N interaction on mycorrhizal colonisation

Pre-germinated seedlings (var. RGT Voilur) were planted in 50 mL Falcon containers filled with attapulgite (Oil Dri UK), and either inoculated with 200 spores of *Rhizophagus irregularis* DAOM 197198 (Agronutrition) per plant or left as non-mycorrhizal controls. Plants were grown under six distinct nutrient treatments with contrasting P (KH_2_PO_4_) and N (KNO_3_) content: (A) 900 μM P, 1000 μM N, (B) 300 μM P, 2000 μM N, (C) 900 μM P, 2000 μM N, (D) 2400 μM P, 2000 μM N, 900 μM P, 5000 μM N, and (F) 2400 μM P, 5000 μM N. Details on the composition of the nutritive solution are provided in Table S5. Plants were harvested at 5 weeks and aboveground biomass was weighed and roots were separated for AMF colonisation assessment.

### Mycorrhizal colonisation

Roots were stained for 2 min at 95°C in a 5 % ink – vinegar solution after a 10% KOH treatment for 10 min at 95°C. Roots were sectioned into 1 cm fragments, mixed and spread across gridded petri dishes. AMF colonisation was assessed visually under a binocular microscope. The presence or absence of mycorrhizal structures was noted at each intersection between a root fragment and a grid line. Mycorrhizal structures, including arbuscules, internal hyphae and vesicles were recorded for a minimum of 100 intersections per sample.

### DNA and RNA Extraction

Frozen root samples were first manually ground in liquid N using a mortar and pestle. **DNA extraction** was performed with 100 mg of root powder using ZymoBIOMICS 96 DNA Kit (Zymo Research) with 50 µl elution volume. Following manufacturer protocol, samples were additionally grinded for 10 minutes using a mixer mill Retsch MM 400. For **RNA extraction**, 100 mg of root powder were transferred in 2 ml tubes containing one metal bead (4mm) / tube, and further grinded twice for 1 minute in a mixer mill Retsch MM 400. Total RNA was extracted using the NucleoSpin® RNA Plus Kit (Macherey Nagel). The quality and integrity of a subset of the RNA samples were assessed using Agilent RNA 6000 Nano Chips (Agilent Technologies), while RNA quantity was measured using a Nanodrop spectrophotometer.

### Analysis of AM marker gene expression

We identified by Blast search the orthologs in wheat of genes coding for the well-conserved ammonium transporter (*SymAMT*), nitrate transporter (*SymNPF*), phosphate transporter (*SymPT*), energising proton pump ATPase (*SymATPase*) involved in the mycorrhizal nutrition (Harrison *et al*., 2002; Wang *et al*., 2014; Breuillin-Sessoms *et al*., 2015; Wang *et al*., 2020) and of a gene (*AM3*) coding for a secreted LysM protein (Yu *et al*., 2023; Tian *et al*., 2024) used as an early AMF colonisation marker gene (Gutjahr *et al*., 2008). These marker genes were used as proxies for symbiotic nutrient transport and the establishment of the AM. Primers were designed, when possible, on conserved regions in the homoeologous (A, B and D) copies in various wheat genomes. The primers used for amplification of the AM marker genes in wheat and the reference genes (*GAPDH, TUB, TEF*) are listed in Table S6. Complementary DNA (cDNA) synthesis was then performed through reverse transcription with the 5X PrimeScript RT Master Mix (Takara), involving an incubation step at 37°C for 15 minutes, followed by 5 seconds at 85°C. qRT-PCR was conducted at an annealing temperature of 60°C using Takyon No ROX SYBR 2X MasterMix blue dTTP (Eurogentec) on a CFX Maestro (Bio-Rad). Gene expression levels were then calculated relative to three wheat housekeeping genes with technical triplicates performed for each biological replicate.

### Sequencing of root-associated fungal communities

ITS2 rRNA amplicon libraries were generated through nested PCR: first, using rcAMDGR (5’-ATGATTAATAGGGATAGTTGGG-3’) (Sato *et al*., 2005) & ITS4 (5’-TCCTSCGCTTATTGATATGC-3’) (White et al., 1990) chosen to exclude wheat ITS2 rRNA, and secondly using ITS 86F (5’-GTGA+ATCATCGAATCTTTGAA-3’) (Turenne *et al*., 1999) & ITS4, which generate an amplicon size compatible with Illumina MiSeq. For both PCR, GoTaq® G2 DNA polymerase (Promega) was employed within the following mixes, PCR 1 on 1µl of DNA extraction: 1X GoTaq® Reaction Buffer, 0.08mM dNTPs, 0.32µM of both primers, 0.04U/µl of polymerase, and 2% DMSO; PCR 2 on 1µl of PCR 1 product: 1X GoTaq® Reaction Buffer, 0.2mM dNTPs, 0.4µM of both primers, 0.02U/µl of polymerase, and 2.5 mM of MgCl2. Nuclease-Free Water for Molecular Biology (Sigma-Aldrich) was used for each PCR mix. PCR were performed with initial denaturation at 95°C for 5 min and final elongation at 72°C for 10 min. PCR 1 entailed 40 cycles at 95°C for 5 sec, 62°C for 30 sec, and 72°C for 1 min. PCR 2 entailed 35 cycles at 95°C for 30 sec, 55°C for 30 sec, and 72°C for 1 min. The resulting ∼300 bp amplicon was visualised via electrophoresis on a 2% agarose gel. The subsequent library creation followed the Illumina two-step PCR protocol and underwent normalisation using SequalPrep plates (Thermofischer). Paired-end sequencing was performed at the Bio-Environment platform (University of Perpignan Via Domitia, Perpignan, France) on a MiSeq system (Illumina) using v2 chemistry with a 2×250 bp read length, following the manufacturer’s protocol.

### Bioinformatics

Raw reads were processed using the pipeline FROGS version 4.1 (Bernard *et al*., 2021) via the Galaxy platform hosted by INRAE Toulouse (https://vm-galaxy-prod.toulouse.inrae.fr/galaxy_main/). This involved a series of steps: merging reads, pre-quality filtering, dereplication, and trimming at 400 bp using VSEARCH version 2.17.0 (Rognes *et al*., 2016), flash version 1.2.11 (Magoč & Salzberg, 2011), and cutadapt version 2.10 (Martin, 2011). Next, clustering was performed through the SWARM algorithm (Mahé *et al*., 2014) and chimeras and singletons were eliminated. A minimum proportion filter of 0.005% of all sequences was applied to retain zero-radius Operational Taxonomic Units (zOTUs) (Escudié *et al*., 2018). An ITSx filter version 1.1.2 (Bengtsson-Palme *et al*., 2013) was employed to identify and extract ITS2. Taxonomic assignment was performed by blastn+ (Camacho *et al*., 2009) using the UNITE Fungi 8.3 database (Kõljalg *et al*., 2013). Clean sequences are deposited at SRA (bioproject ID PRJNA1291988 available at: http://www.ncbi.nlm.nih.gov/bioproject/1291988).

### Data Analysis

All statistical analyses were performed using R 4.1 using the packages detailed in each section.

#### Effects of treatments on mycorrhizal functioning and mycorrhizal colonisation

The effects of the experimental treatments on the expression of the AM marker genes (field & experiment 2), were analysed by ANOVA with a posterior Tukey test for multiple comparisons. ANOVA assumptions were evaluated using the R package ‘performance’. In the case a response variable violated assumptions of homogeneity of variances and/or residual normality, we performed a non-parametric Kruskal-Wallis test followed by a Dunn test for multiple comparisons with Benjamini-Hochberg correction (R package ‘dunn.test’). For experiment 3, a linear model was fitted to assess the interaction of N and P availability on root colonisation by AMF. The significant interaction was visualised using the ‘interactions’ R package.

#### Analyses of fungal communities in wheat roots

*Datasets -* Metabarcoding data of root-associated fungal communities were processed with the R ‘phyloseq’ package (McMurdie & Holmes, 2013). We examined rarefaction curves for each sample to assess sequencing quality. Next, we merged biological replicates per plot in order to obtain the fungal communities for each experimental plot. Rarefaction curves generated for each plot can be found in Fig. S1. To evaluate changes of AMF communities (belonging to the Glomeromycota phylum) with P fertilisation treatments, we investigated community composition in terms of proportion of different taxa, indicator species analysis, alpha diversity, and we examined community structure with ordination and PERMANOVA analyses. Due to the issues associated with using rarefied or relative abundance data for community structure analyses of compositional data (McMurdie & Holmes, 2014; Gloor *et al*., 2017), different approaches or data transformations were applied for each analysis (specified below).

##### Community composition

zOTUs belonging to the Glomeromycota phylum were extracted from the fungal dataset and relative abundances at Genus level were calculated and plotted in compositional plots according to P treatment and sampling year. Unclassified and multi-affiliated taxa at genus level were merged into an unidentified Genus group. To explore the contribution of specific zOTUs to community composition in each P treatment, we plotted rank abundance curves (R package ‘BiodiversityR’) and the 20 most abundant zOTUs (Glomeromycota phylum, CLR normalised) into a heatmap (R package ‘microbiomeutilities’).

##### Indicator species analysis

To explore preferential zOTU associations to P treatments, we used indicator species analysis (De Cáceres & Legendre, 2009), which calculates an association index (IndVal) between taxa and treatment, and then tests for its significance with a permutation test (R package ‘indicspecies’). IndVal is a product of two components: A which corresponds to an estimate of the probability that a sample belongs to a treatment (specificity), and B which is an estimate of the probability finding a taxon in a treatment (fidelity). These components give additional information on why a taxon can be used as an indicator.

##### Alpha diversity

Alpha diversity of AMF communities (Richness, Shannon index, Gini-Simpson index, Pielou’s evenness and dominance index) was estimated per P treatment (R package ‘microbiome’). To calculate alpha diversity indices, raw data were rarefied to equal number of fungal reads (minimal sample reads: 20.704 reads in 2019 and 34.957 in 2022). Within a year, variation in alpha diversity indices across P treatments was evaluated using Kruskal-Wallis tests followed by a Dunn test for multiple comparisons with Benjamini-Hochberg correction.

##### Beta diversity

A permutational multivariate analysis of variance (PERMANOVA, R package ‘vegan’) was then performed on Aitchison distances (CLR transformed AMF raw data, R package ‘microbiome’) to test the effects of the P treatments and the experimental block on AMF community structure. Multiple pairwise comparisons were performed for significant PERMANOVA results and were corrected via the Bonferroni correction (R package ‘pairwise.adonis’). All significant PERMANOVA results were further analysed with permutational multivariate analyses of dispersion (PERMDISP, R package ‘vegan’) to verify that all significant PERMANOVA results represented a difference in average community composition, rather than a difference in dispersion of communities among treatments. To visualise differences in AMF community composition, Aitchison distances were ordinated using principal components analysis (R package ‘PCAtools’).

## Results

### Validation of the arbuscular mycorrhiza (AM) marker genes in controlled conditions

In order to estimate the mycorrhizal nutrition in field-grown wheat, we designed primers to measure the expression of well-conserved genes coding for an ammonium transporter (thereafter called *SymAMT*), a nitrate transporter (thereafter called *SymNPF*), and a phosphate transporter (thereafter called *SymPT*), as well as an energising proton pump ATPase (thereafter called *SymATPase*), specifically expressed in arbuscule-containing cells and known to be localised in the peri-arbuscular membrane. We tested the specificity of the primers and of the gene expression pattern by measuring their expression in mycorrhizal and non-mycorrhizal roots of wheat grown under controlled conditions.

All genes were only detected in plants inoculated with *R. irregularis* (Fig. 1). In addition, in order to estimate AMF colonisation at the same time as the mycorrhizal nutrition, we designed primers against a gene (*AM3*) coding for a secreted LysM protein specifically expressed in arbuscule-containing cells and used as an early AMF colonisation marker on other species. Again, this gene expression was only detected in mycorrhizal roots (Fig. 1).

**Fig. 1.**
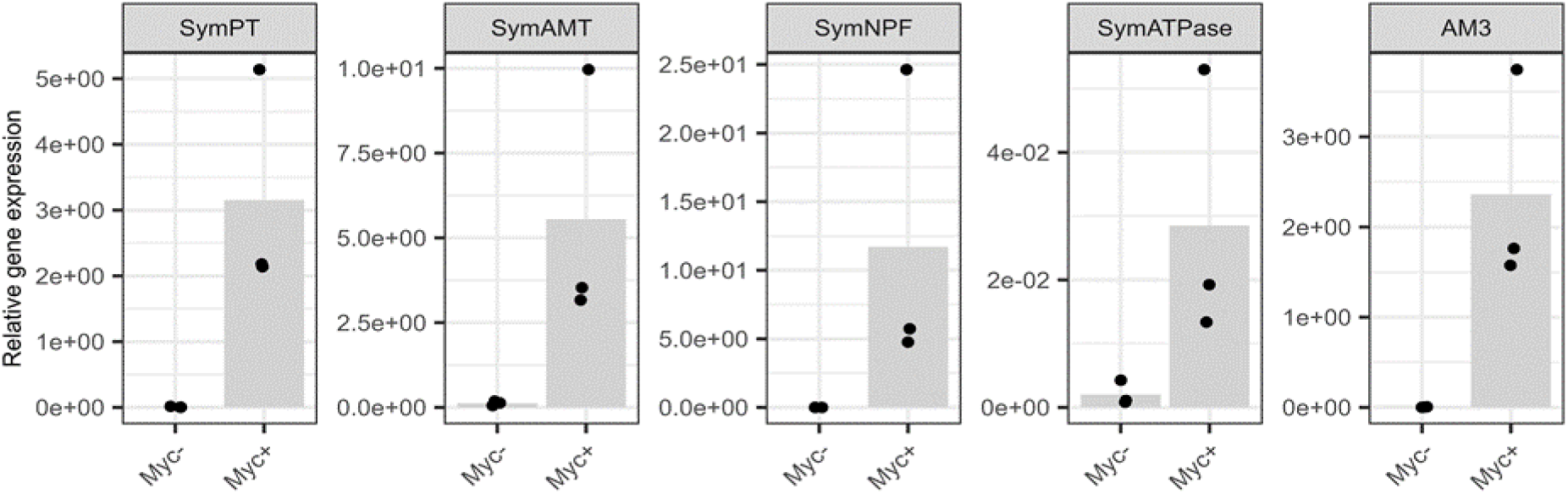
Selected AM marker genes were specifically expressed in mycorrhizal wheat roots. Relative expression of genes coding for the peri-arbuscular nutrient transporters/pump (SymPT, SymAMT, SymNPF, SymATPase) and of an AMF colonisation marker gene (AM3) measured by RT-qPCR in mycorrhizal (inoculated with R. irregularis, Myc+) and non-mycorrhizal (control, Myc-) roots of wheat grown on attapulgite under controlled conditions (n= 3)). Bars represent the mean, individual plant pools are shown as black dots.

### Nutrient transporter expression and N availability in field conditions

We then measured the expression of the nutrient transporters and of the colonisation marker gene in the roots of wheat grown in field plots that received various levels of long-term P fertilisation (No P, Low P and High P). Patterns of relative gene expression according to P treatments differed between years (Fig. 2). While in 2022 there was a pattern of decreased relative gene expression of AM marker genes as P fertilisation increased, this was not observed in 2019. In contrast, the expression of the symbiotic ammonium transporter (*SymAMT*) increased with increased P availability in 2019.

**Fig. 2.**
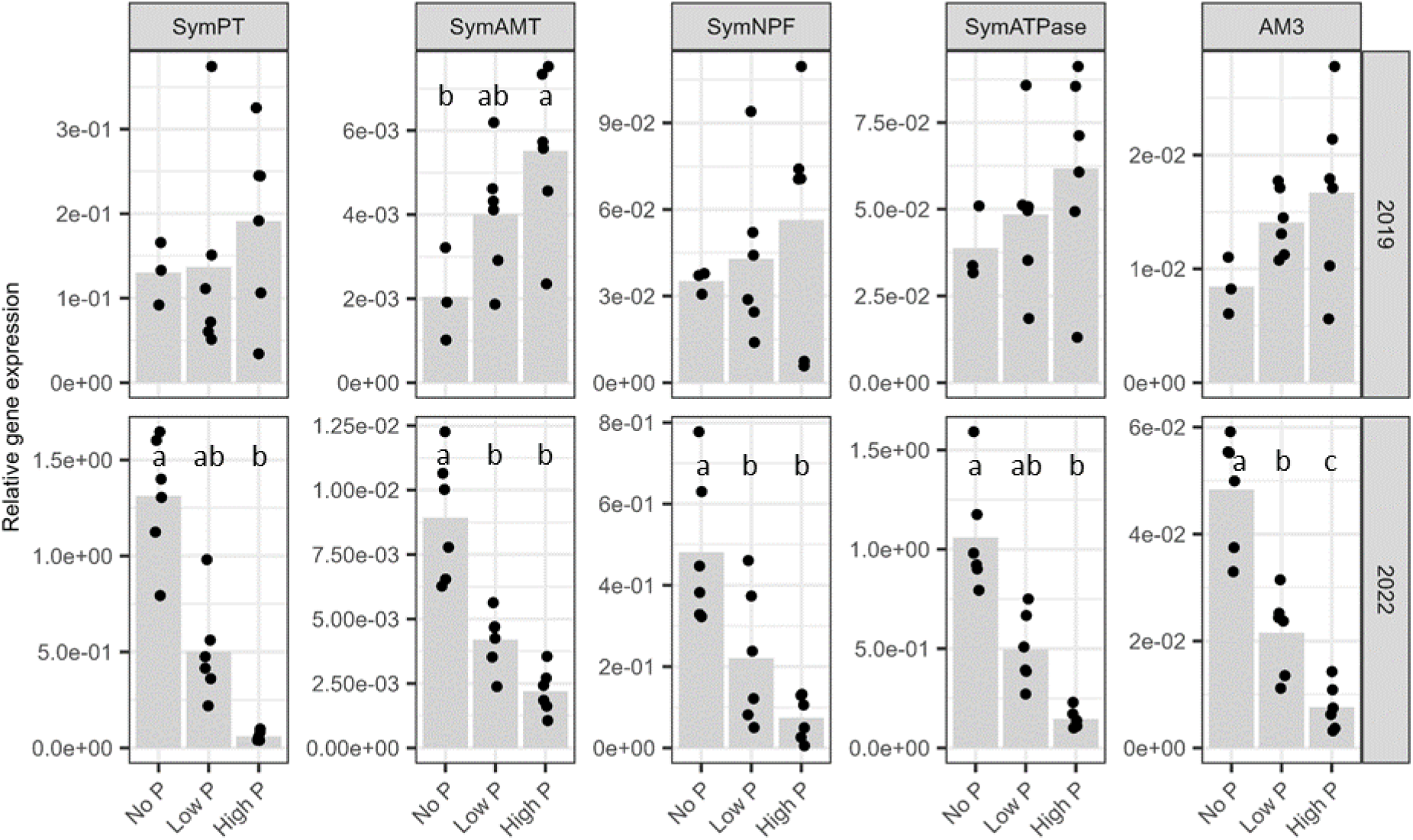
P fertilisation inhibited AM in wheat roots sampled in 2022 but not in those sampled in 2019. Relative expression of AM marker genes in wheat roots measured by RT-qPCR sampled in a long-term P fertilisation trial (treatments No P, Low P or High P) in 2019 or 2022 (n= 3 - 6). Bars represent the mean, individual plant pools are shown as black dots. Different letters indicate significant differences in AM marker gene expression among P treatments (p-value <0.05, detailed in Table S7) assessed by ANOVA followed by post hoc Tukey test or by Kruskal-Wallis test followed by Dunn test with Benjamini-Hochberg correction when ANOVA assumptions were not met. Note the difference of scale among AM marker genes and years.

We hypothesised that the effect of the years could be related to variability in the level of N availability at the sampling timepoint. To test it, we measured wheat root N content (%N), as a proxy of N availability, on the same root material used for RT-qPCR. Root N content differed between years with N (%) 1.5-fold greater in 2022 than in 2019 (Fig. 3). In 2019, %N content differed between P treatments with high P roots presenting lower N than the No P treatment. No significant differences in root %N were detected among P treatments in 2022, although a similar trend was observed.

**Fig. 3.**
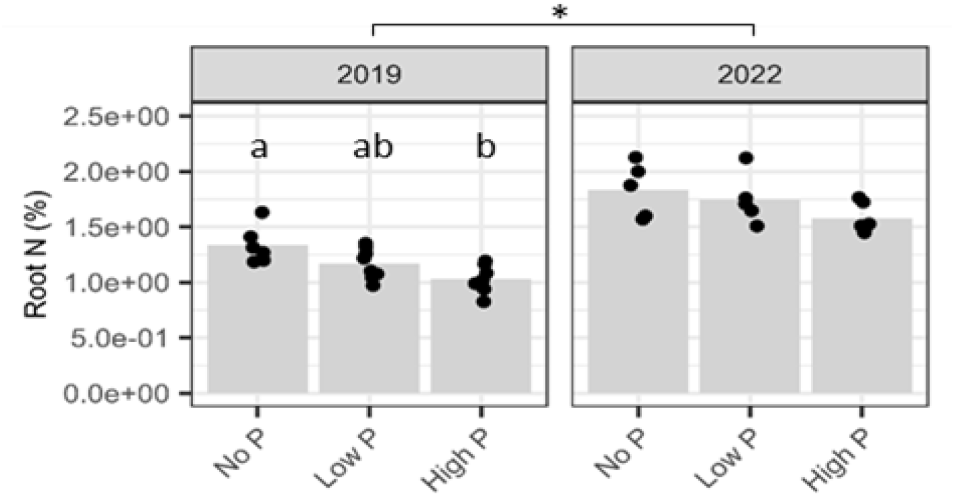
Root N content (%) in field conditions was greater in 2022 than in 2019 (n= 5 to 8). Bars represent the mean, individual plant pools are shown as black dots. Different letters indicate significant differences (p-value <0.05, detailed in Table S8) assessed by ANOVA followed by post hoc Tukey test. Asterisk shows a difference (p-value <0.05, detailed in Table S8) between sampling years for all P treatments combined using ANOVA.

### Effects of N and P availability on the AM interaction

To confirm that interaction between N and P availability influences AM functioning in wheat, soils collected from the long-term P field trial were used in combination with a N fertilisation treatment in a controlled condition experiment (*experiment 2*). AMF colonisation, measured by the expression of the marker gene *AM3* (Fig. 4a) and by microscopy (Fig. 4b), as well as the mycorrhizal nutrition (expression of the AM-specific nutrient transporters, Fig. 4a) responded to the interaction between N and P availability in a similar manner as in field conditions. No difference in AMF colonisation was observed in the experimental treatments in which one of the nutrients was limiting, (i) *No P*, limited availability of both P and N, (ii) *No P+N*, limited availability of P but sufficient N availability, and (iii) *High P*, limited N availability but sufficient P availability. In contrast, AMF colonisation was inhibited in the *High P+N* treatment, where both N and P were sufficient. Interestingly, *SymNPF* and *SymATPase* expressions were upregulated by high P treatment only when N was limiting, while *SymPT* was upregulated in all treatments where P was limiting. In sum, wheat plants showed increased AMF colonisation in treatments with a limiting nutrient, yet N and P transporters showed contrasting expression regulation depending on the identity of the limiting macronutrient.

**Fig. 4.**
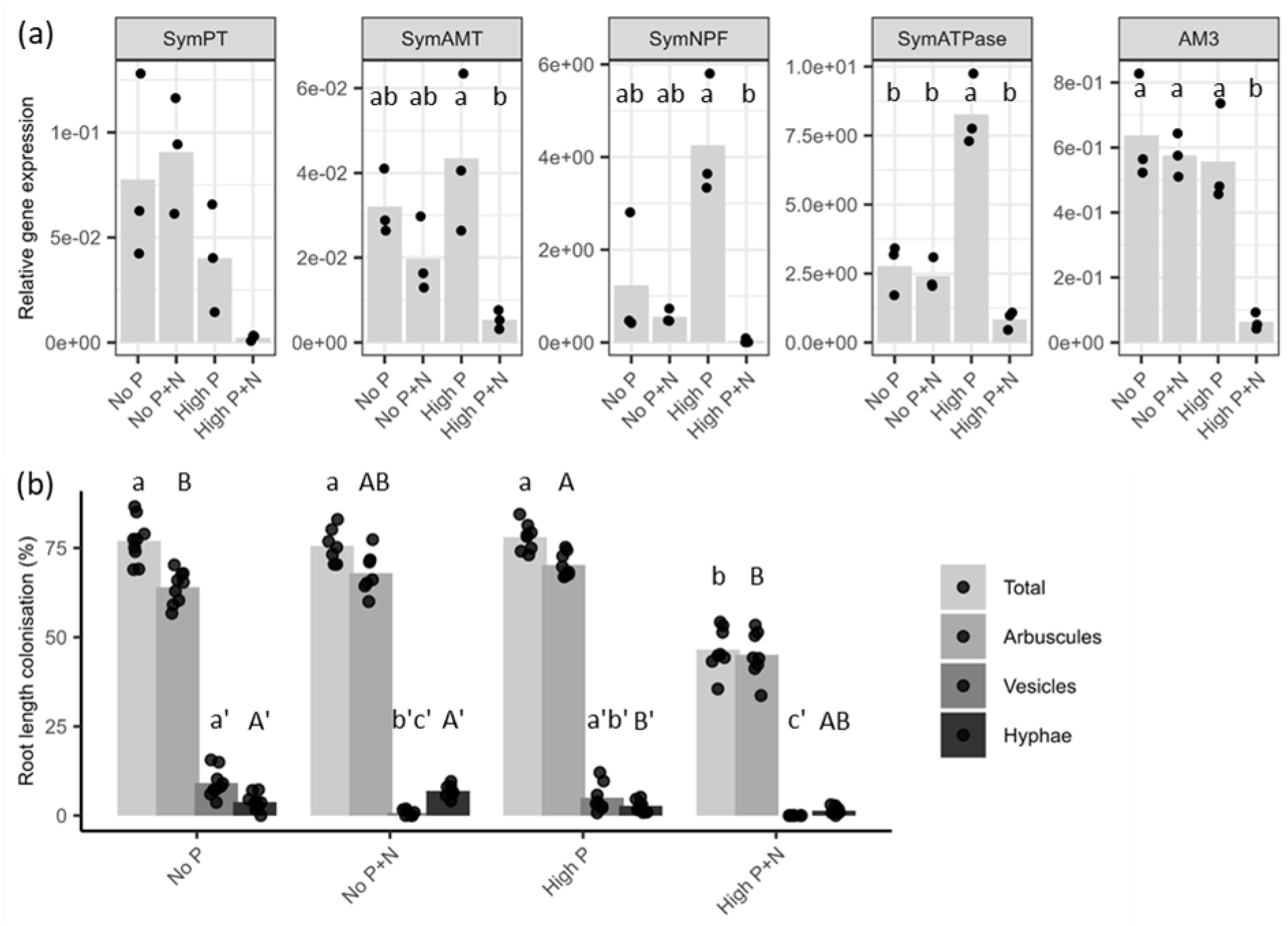
Expression of the mycorrhizal nutrition genes is modulated by soil N and P availability in wheat plants grown in controlled conditions. (a) Relative expression of AM marker genes was measured by RT-qPCR on wheat plants grown under controlled conditions on soils with distinct P availability (soils from long-term P fertilisation field trial No P or High P) and N fertilisation treatment (+N corresponds to fertilisation 2000 µM of nitrate) (n= 3). Bars represent the mean, individual plant pools are shown as black dots. Different letters indicate significant differences (p-value <0.05, detailed in Table S9) in the AM marker gene expression among P treatments assessed by ANOVA followed by post hoc Tukey test or by non-parametric Kruskal-Wallis test followed by Dunn test with Bonferroni correction when ANOVA assumptions were not met. (b) AMF root length colonisation was measured using the grid intersect method in wheat plants grown as in A. Bars represent the mean, individual plants are shown as black dots (n= 7-9). Different letters indicate significant differences (p-value <0.05, detailed in Table S9) using Kruskal-Wallis test followed by post-hoc Dunn test with Bonferroni correction.

A second controlled experiment (*experiment 3*) was performed to study in more detail the interaction of N and P availability in the soil solution (soil) on root AMF colonisation. As in *experiment 2*, results show an increase of AMF colonisation when one macronutrient was limiting (Fig. 5b; considering treatment F as no limitation). A linear model further confirmed that the interaction between P and N availability explained mycorrhizal colonisation of wheat roots (Fig. S2). Biomass, in turn, was greatly determined by N availability (Fig. 5c, treatments A-C-E & D-F) but not by P availability (Fig. 5c, treatments B-C-D & E-F).

**Fig. 5.**
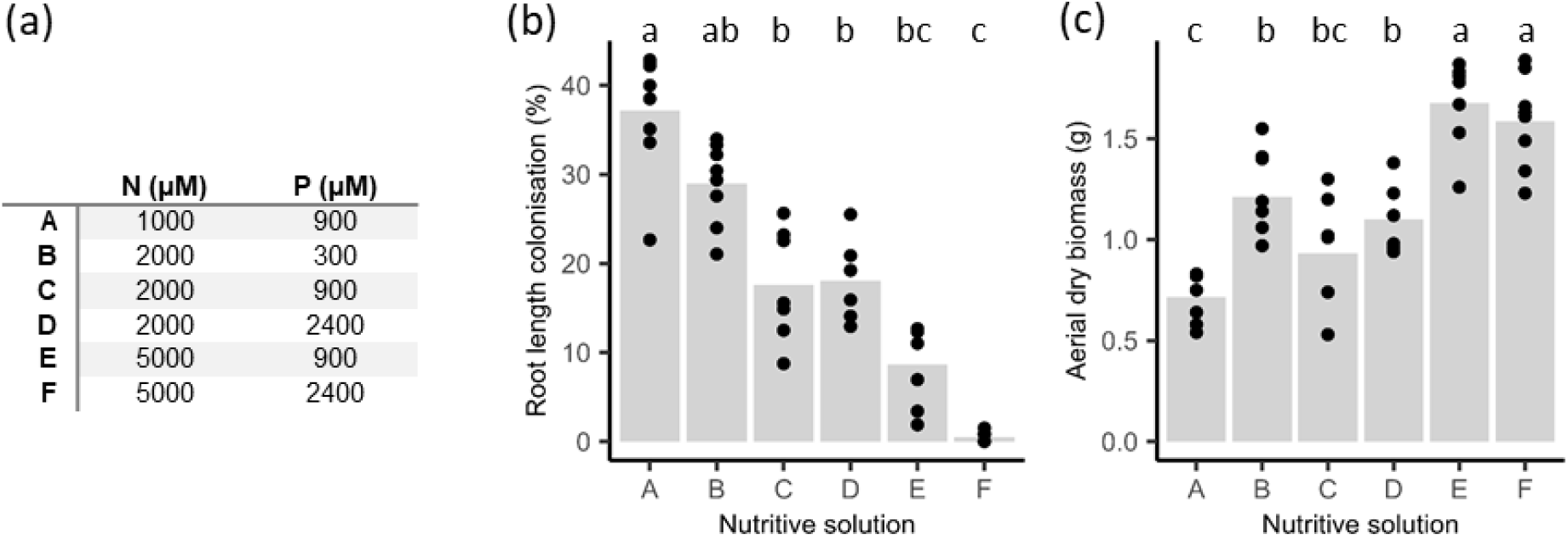
Limiting N and P enhances wheat root AMF colonisation by R. irregularis in artificial soil. Wheat plants were grown under controlled conditions in attapulgite using a range of 6 nutritional treatments with a gradient in N and P concentration. Wheat was grown for 5 weeks after inoculation with 200 spores of R. irregularis DAOM197198. (a) Details of the 6 nutrient conditions tested in terms of N (nitrate) and P concentration. (b-c) Above-ground dry biomass (b) and AMF root length colonisation measured using the grid intersect method Different (c). Bars represent the mean, individual plants are shown as black dots (n = 7-9). Different letters indicate significant differences (p-value <0.05, detailed in Table S10) assessed by ANOVA followed by post hoc Tukey test (b), and Kruskal-Wallis test followed by post hoc Dunn test with Benjamini-Hochberg correction (c).

### AMF communities in wheat roots in a long-term P fertilisation assay

In addition to investigating the effect of P treatments on AMF colonisation and mycorrhizal nutrition, we analysed its effect on root AMF communities by metabarcoding. After bioinformatic processing, 654.338 and 820.217 sequences obtained by Illumina sequencing of an ITS2 amplicon were retained and clustered into 580 and 738 zOTUs for 2019 and 2022, respectively. A total of 573 and 722 zOTUs were identified as fungi using the UNITE database (with zOTUs not classified as fungi representing 3.60 and 8.83% of the reads). From this set, 524 zOTUs (2019) and 643 zOTUs (2022) were assigned to a specific fungal phylum. The phylum *Glomeromycota, i*.*e*. AMF, comprised 79 zOTUs in 2019 and 148 zOTUs in 2022, representing 4.66 % and 6.31% of the total reads, respectively, with similar proportions across P treatments (Fig. S3). Although beyong the scope of this study, other fungal phyla increased their relative abundance with increased P fertilisation. The relative abundance of the *Olpidiomycota* increased with P fertilisation in both years, while an increase in *Mortierellomycota* relative abundance with P fertilisation was observed only for 2022. Taxonomic affiliation and zOTU tables are provided in Tables S11-S14.

AMF fungal communities in wheat roots were dominated by zOTUs belonging to the family *Glomeraceae* for all treatments and years (Fig. 6). There was a trend of increased relative abundance of the genus *Funneliformis* with long-term P fertilisation for both campaigns, and a complementary trend of decreased proportion of zOTUs belonging to non-identified *Glomeraceae*, with greater differences in relative abundance of *Funneliformis* genus between P treatments for the year 2022.

**Fig. 6.**
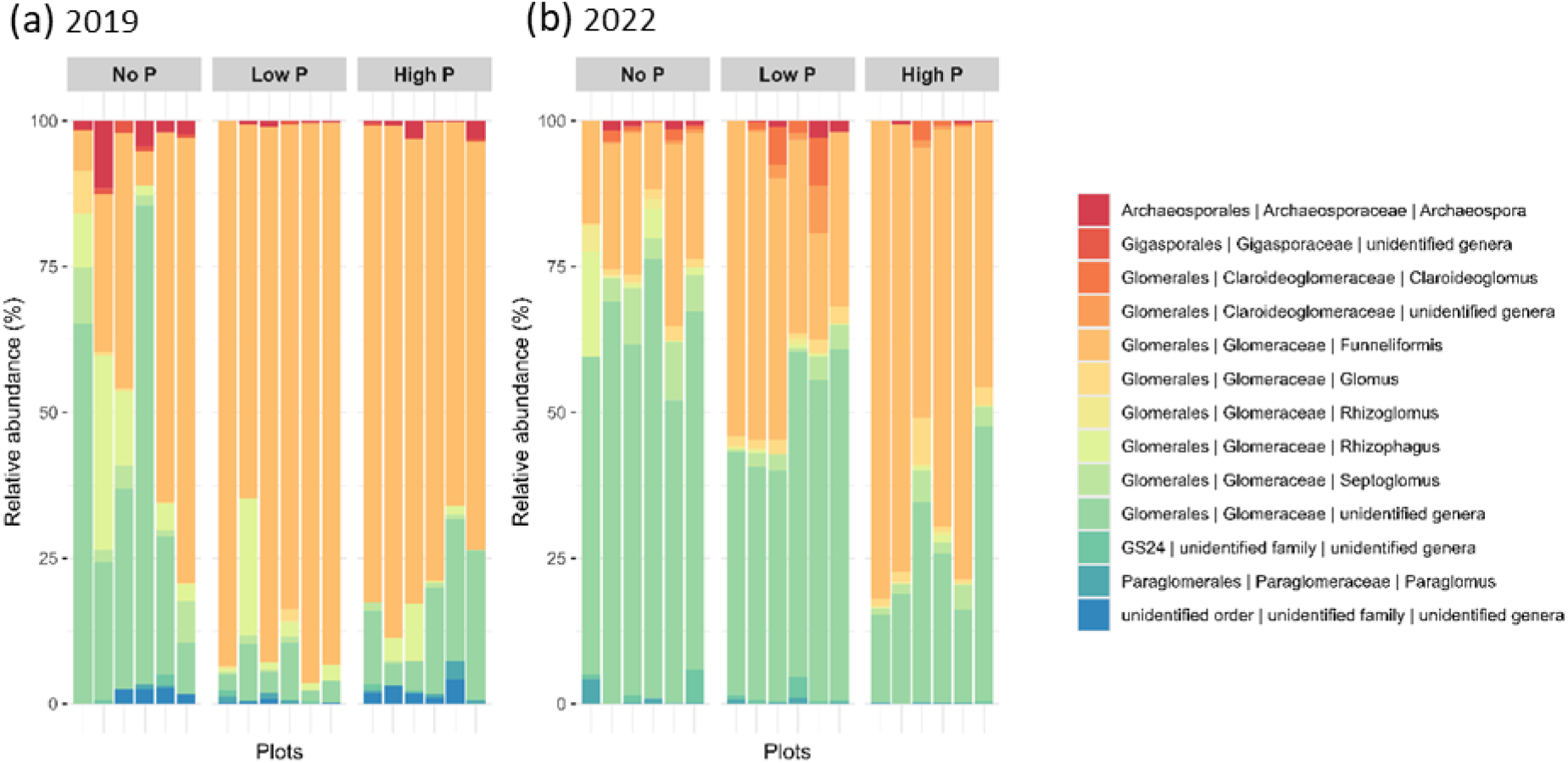
Relative abundance of AMF genera in wheat roots from a long-term P fertilisation field trial. P fertilisation treatments: No P, Low P, and High P. Wheat was sampled in (a) 2019 and (b) 2022. Relative abundances of AMF were calculated based on Illumina sequencing of an ITS2 amplicon.

Similar patterns were observed when AMF composition was evaluated at the zOTU level. The most dominant zOTUs in 2019 were affiliated as *Funneliformis mossae* [Cluster_126, Cluster_132 & Cluster_91] in Low P and High P treatments with the most abundant cluster [Cluster 126] presenting 15% of the total number of reads, while for the No P treatment the most abundant cluster [Cluster_94], also *F. mossae*, represented 8% of the reads. In 2022, the *Funneliformis mossae* [Cluster_99] dominated communities in all P treatments but with increasing relative abundance as P fertilisation increased (9% for No P, 16 % for Low P and 24% for High P). Rank abundance curves showing AMF zOTU dominance and heatmaps showing the distribution of the most abundant AMF zOTUs among plots and treatments can be found in Fig. S5 and S6.

The patterns of root-associated AMF communities alpha diversity indices at the zOTU level differed between years. In 2019 there were no differences in alpha diversity indices across P fertilisation treatments. In contrast, in 2022, alpha diversity indices Gini-Simpson and Pielou’s evenness decreased as P availability increased (Fig. 7, differences between P fertilisation extremes). Community dominance was the greatest under the high P treatment, which was on average 2-fold greater than the No P treatment (Fig. 7). In sum, in 2022, plants grown in High P treatment soils presented less diverse communities which were dominated by a smaller number of zOTUs in comparison with the No P treatment (Fig. S5 and S6).

**Fig. 7.**
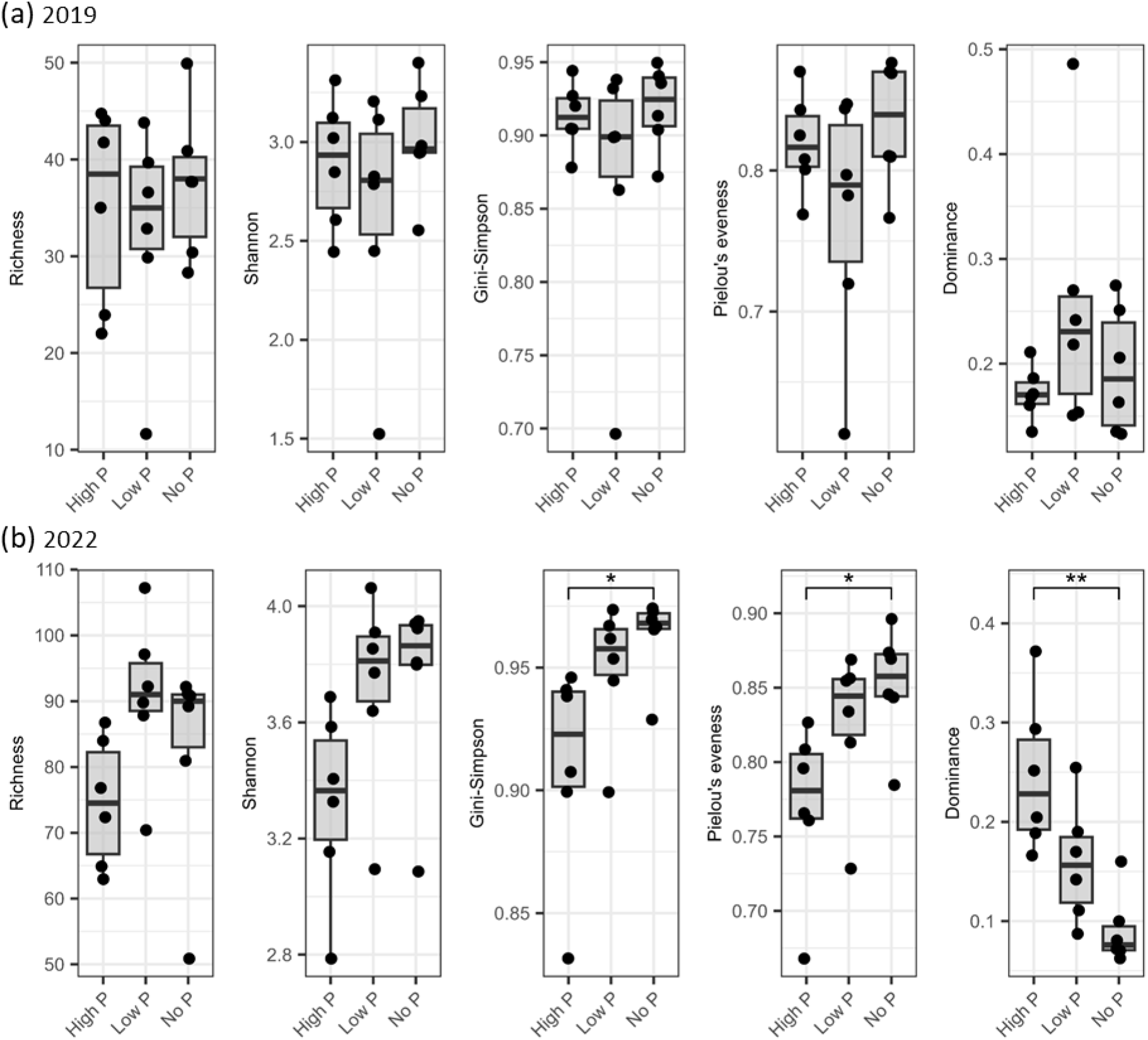
Alpha diversity metrics of AMF zOTUs in wheat roots from a long-term P fertilisation field trial. Wheat plants were sampled in (a) 2019 and (b) 2022. Differences among treatments were assessed by Kruskal-Wallis test followed by post hoc Dunn test with Benjamini-Hochberg correction. Asterisks show significant differences among treatments (p<0.05, detailed in Table S18).

An indicator species analysis showed that a number of zOTUs belonging to the *Funneliformis* genus (mostly *F. mossae*) were indicators of P fertilisation in 2019 (11 zOTUs) and 2022 (9 zOTUs) (Fig. 8). In contrast, zOTUs belonging to the *Glomus* genus were indicators of both No P and Low P treatments in 2022, while few *Septoglomus* and unidentified *Glomeraceae* genus zOTUs were associated with No P in both years. The identity indicator zOTUs per P treatment and associated IndVal values are available in Tables S15 and S16 for 2019 and 2022, respectively.

**Fig. 8.**
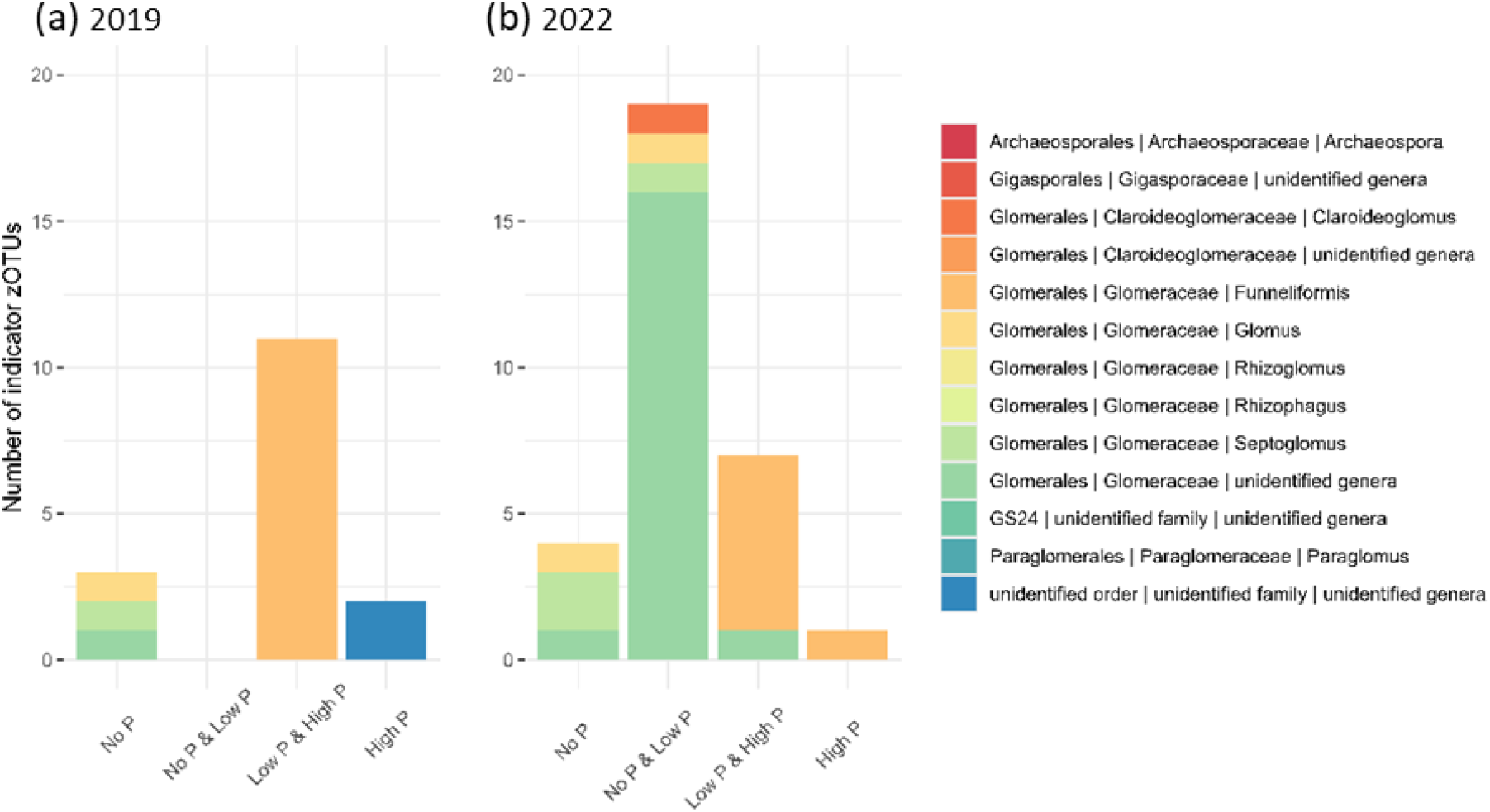
Number of AMF zOTUs which are indicator taxa closely associated with either No P, No P & Low P, Low P & High P or High P treatments in wheat roots from the long-term P fertilisation field trial. Wheat plants were sampled in (a) 2019 and (b) 2022.

In addition, the structure of AMF communities at the zOTU level differed between P treatments for both years (Fig. S6). Pairwise contrasts revealed differences among No P and High P treatment after adjusting for multiple comparisons but not according to experimental block (Table S17). Differences in multivariate dispersions among P treatments were detected for year 2019, with higher variation in community structure for the No P treatment in comparison to the P addition treatments (Fig. S6). The visualisation of community structure by PCA showed that P treatments separated by the PC1 axis with 27.5 and 29.9 % of variation explained in 2019 and 2022, respectively (Fig. S6).

## Discussion

In this study we investigated the effects of long-term P fertilisation on root-associated AMF communities in wheat and their role in nutrient acquisition under field conditions. It should be noted that, in this field experimental system, the gradient of availability of P is strong, ranging from very limited (No P) to an excess (High P) of P for crop development. This is reflected in the wheat yields during the two campaigns (Table S4) and in the yields of other crops previously grown in this field trial (Nawara *et al*., 2017). We measured the expression of peri-arbuscular transporters/pump involved in nutrient uptake and characterised the AMF community in roots of wheat grown in plots with varying P availability over two distinct years. We hypothesised that (i) the mycorrhizal nutrition is modulated by nutrient availability, and that the expression of AM transporters will follow a similar pattern; and that (ii) changing soil P availability will change AMF diversity shifting community composition towards species that better tolerate or compete in each environment. The results show that the composition and structure of root-associated AMF communities differed according to long-term P treatment and that the strength of differences varied depending on the year of sampling. Likewise, the differences in AM transporter expression among treatments was dependent on the year. The distinct N content in the roots at the time of sampling led us to hypothesise that the differences observed between the different years were likely to be the result of the interaction between the availability of P and that of the other key macronutrient, N. The interaction between P and N for AM colonisation and functioning is well acknowledged in the literature (Sylvia & Neal, 1990; Johnson, 2010; Javot *et al*., 2011; Bonneau *et al*., 2013; Nouri *et al*., 2014; Püschel *et al*., 2016; Corrêa *et al*., 2024) and was further supported by experiments under controlled conditions. Finally, we showed that the expression of selected wheat AM-marker genes involved in arbuscule formation and plant nutrient uptake can be used to explore the contribution of the AMF community to plant nutrient acquisition.

Our results suggest that the interaction between the availability of the two major macronutrients, N and P, can partly explain the between-year differences in AM functioning in wheat in the same long-term P fertilisation trial. Although similar N fertilisation was applied in 2019 and 2022, the wheat N nutrition status differed between the campaigns. This could be due to interactions between N fertilisation and meteorology, or crop succession (sunflower and soybean were grown before wheat in 2019 and 2022, respectively), which can dramatically affect N availability in soil. Alternatively, this could also be due to higher wheat N demand in 2019. Indeed, yield was 40% greater in 2019 than in 2022 for all P treatments (Table S4). Furthermore, the crop nutritional requirements change throughout the season influencing the formation and maintenance of mycorrhizae and their functioning. The formation and degeneration of arbuscules is highly dynamic (Kobae & Hata, 2010; Kobae *et al*., 2016; Gao *et al*., 2019), allowing an adjustment of the AM to changes in plant nutritional status (e.g. nutrient limitation/starvation) and the soil nutritional context. By measuring expression of genes encoding peri-arbuscular transporters/pump, we aimed to obtain a snapshot into the nutritional functions provided by AMF to their host plant at a given point in the season. While in 2022 we observed inhibition of the AMF colonisation following increased soil P availability, in line with previously published data on the same field trial (Tang *et al*., 2016), and consequently a decrease in the expression of the AM transporters/pump, we did not observe such responses in 2019. Our results suggest that the wheat N status greatly modulated mycorrhizal nutrition under field conditions. In 2019, N limitation may have reduced the need for enhanced P uptake in wheat plants under limiting P availability, while increasing the need for N uptake under non-limiting P availability. This aligns with previous work on maize showing that the activation of the transcriptional response to P starvation is contingent on N availability (Schlüter *et al*., 2012; Torres-Rodríguez *et al*., 2021), with the N limitation response dominating the transcriptome when plants were subjected to joint N and P limitation.

The controlled condition experiment using soil from the field trial confirmed that the expression of peri-arbuscular transporters regulated in wheat roots depends on the identity of the limiting macronutrient for the plant. P limiting conditions, no matter the N fertilisation treatment, promoted the expression of the phosphate transporter (*SymPT*), while in P sufficient and limiting N conditions the symbiotic nitrate transporter (*SymNPF*) was upregulated. These regulations are independent of the level of colonisation since neither the expression of the *AM3* marker gene nor the microscopy quantification changed between N-limiting/P-sufficient and N-sufficient/P-limiting conditions, while expression of *SymPT, SymAMT* and *SymNPF* changed drastically between these conditions. Interestingly, we found that *SymAMT* was upregulated in N-limiting/P-sufficient conditions in the field in 2019, while *SymNPF* was upregulated in controlled conditions under the same conditions. This difference in transporter expression may be a consequence of distinct N forms used as fertiliser (nitrate in controlled conditions and ammonium nitrate in the field) and suggests that the regulation of peri-arbuscular N transporters is specific to the nature of N available in the soil. Altogether, these results also demonstrate that despite being commonly used as markers of AMF colonisation, the expression of peri-arbuscular transporters is not a reliable indicator of colonisation. Conversely, the expression of the *AM3* marker gene in roots followed the same trend as the level of AMF colonisation quantified by microscopy under both controlled conditions and in the field trial (Fig. 4 and S7), suggesting that the expression of the *AM3* marker gene is a reliable indicator of AMF colonisation. The expression of the AM3 at a low colonisation level (when nutrition may not be the main benefit of AM) remains to be tested. Globally, our results support the idea that the use of these AM marker genes and transporters can be used as a proxy of mycorrhizal functioning of the AM at a moment in time when comparing treatments with contrasting nutrient availability. Next steps will be to link the transporter expression with measures of plant nutrient acquisition via the mycorrhizal pathway by means of isotopic tracer experiments.

Our hypothesis that a long-term increase of soil P would lead to a reduction in AMF diversity and a shift in AMF community composition towards species adapted to high-P environments in wheat roots was partly validated. We found that P fertilisation affects the composition and structure of the wheat-associated AMF community but not its richness, as already described for other crops in previous field trials (Beauregard *et al*., 2010; Lang *et al*., 2018; Dueñas *et al*., 2020). The majority of AMF zOTUs in wheat roots belonged to the *Glomeraceae* family in all P treatments. *Glomeraceae* in general exhibit ruderal life-history traits according to Grime’s C-S-R framework (fast growth rates and preferential allocation of biomass to roots rather than to extraradical hyphae), which are associated with conditions of high disturbance and low stress, and are thus dominant in agricultural ecosystems (Oehl *et al*., 2003; Ujvári *et al*., 2023). Within the *Glomeraceae*, we observed a consistent shift in the root AMF community composition for both sampling years, with zOTUs of the *Funneliformis* genus gaining prevalence with increased P availability. The increased prevalence of the *Funneliformis* genus with P fertilisation has previously been described in roots of maize (Lang *et al*., 2018) and wheat (Peng *et al*., 2024). A similar correlation had also been observed between the relative abundance of the *Funneliformis* genus in roots and soil P availability throughout the wheat growth cycle (Bainard *et al*., 2014). Interestingly, the authors showed that while all soil AMF exhibited negative correlations as P levels increased, certain *Funneliformis* OTUs in the roots exhibited positive correlations. They proposed that these OTUs may shift their growth strategy with P availability, potentially investing less in extraradical hyphae and more in root structures. In a long-term fertilisation trial (Williams *et al*., 2017) reported that soil AMF communities remained unaffected in terms of composition with increased soil P, yet AMF reduced their investment in extraradical biomass and attributed such change to either a change in growth habits or decreased carbon allocation to the fungi. Such changes in biomass allocation of certain OTUs (e.g. *Funneliformis*) with fertilisation could explain our results. Another possible explanation could be that certain *Funneliformis* zOTUs are more efficient at the acquisition of other nutrients (e.g. N) that become limiting with increased P fertilisation or that they provide other non-nutritional benefits to plants(Frew *et al*., 2023). Several studies on isolated strains have indeed shown functional variability between species and genotypes, including for ability to provide N or P to their host, supporting a possible shift in the AMF community associated with nutritional functionality (Munkvold *et al*., 2004; Avio *et al*., 2006; Mensah *et al*., 2015).

It is well established that the host plays a crucial role in determining their associated AMF community. Various studies have shown contrasting patterns in AMF community composition in roots and soil in both natural and agricultural ecosystems (Qin *et al*., 2020), increasing the challenge in comparing findings across studies on distinct plant species and potentially distinct nutritional statuses (Johnson, 2010). Interestingly, the strongest differences in root-associated AMF community structure and diversity among long-term P treatments were observed for year 2022, when N was non-limiting to wheat plants (based on %N in wheat roots). In 2022, a small number of unidentified *Glomeraceae* zOTUs were most abundant in No P and Low P treatments compared to High P, and indicators of No P and Low P treatments. These zOTUs might be more adapted at P mobilisation than dominant *Funneliformis mossae* zOTUS which are ubiquitous but dominate in the High P treatment. In contrast, N limitation in soil could increase the competition between plant and fungus for N (Kuzyakov & Xu, 2013; Püschel *et al*., 2016). However, the question remains whether the most abundant zOTUs in our treatments predominate due to their lifestyle traits or due to host filtering and preferential allocation for resources.

A major goal in mycorrhizal research is to understand how AMF diversity and community composition determine host nutrient acquisition. A critical way to achieve that will be to establish a link between AMF communities and AM nutritional functions (Gamper *et al*., 2010; Powell & Rillig, 2018). The present work performed on wheat, a major crop, shows the potential to achieve that by enriching soil with N and P, and combining characterisation of the AMF community composition with the expression of the AM functional marker gene.

## Acknowledgments

We thank Nadine Bouzari for technical help. This study was developped within the framework of the “Laboratoires d’Excellences (LABEX)” TULIP (ANR 10 LABX 41) and of the “Ecole Universitaire de Recherche (EUR)” TULIP GS (ANR 18 EURE 0019). MT PhD was co-funded by the Université Fédérale Toulouse Midi-Pyrénées (UFTMiP) and the Région Occitanie.

## Author contributions

MT, CJ, CR, and BL designed the experiments. CJ, PB and EL performed the field assays and analysis. VG and MT performed RT-qPCR. MT performed all the others experiments, MT and AA analysed the data. MT, BL and AA wrote the manuscript.

## Supporting Information

Fig S1. Rarefaction curves of experimental plots per P treatment.

Fig S2. Interaction graph N X P

Fig S3. Composition of root-associated fungal communities at Phylum level per P treatment and year.

Fig S4. AMF community rank accumulation curves.

Fig S5. Heatmap and cluster analysis AMF zOTUS

Fig S6. PCA representing community structure of AMFs based on Aitchison distances

FigS7. Correlation between the relative expression of the AM3 marker gene and the AMF colonisation level quantified by microscopy.

Table S1. Soil properties measured in the topsoil (0–30 cm depth) along the long-term P fertilisation trial.

Table S2. Sampling date and details on the technical itineraries for the two-durum wheat culture seasons 2018–2019 and 2021–2022.

Table S3. Climatic data for seasons 2018–2019 and 2021–2022.

Table S4. Harvest data for seasons 2018–2019 and 2021–2022.

Table S5. Composition of nutrient solutions used in experiment 1 & experiment 3.

Table S6. Description of the symbiotic markers used in this study.

Table S7. Statistics of the Fig. 2

Table S8 Statistics of the Fig. 3

Table S9: Statistics of the Fig. 4

Table S10: Statistics of the Fig. 5

Table S11: Taxonomy table 2019

Table S12. Taxonomy table 2022

Table S13. zOTU table 2019

Table S14. zOTU table 2022

Table S15. AMF zOTU indicators of P treatments 2019 (indicator analysis results)

Table S16. AMF zOTU indicators of P treatments 2022 (indicator analysis results)

Table S17. Pairwise PERMANOVA test 2019 and 2022.

Table S18: Statistics of the Fig. 7

