## supplementary figures for "Soil phosphate availability modulates the arbuscular mycorrhizal fungal community and mycorrhizal nutrition in wheat"

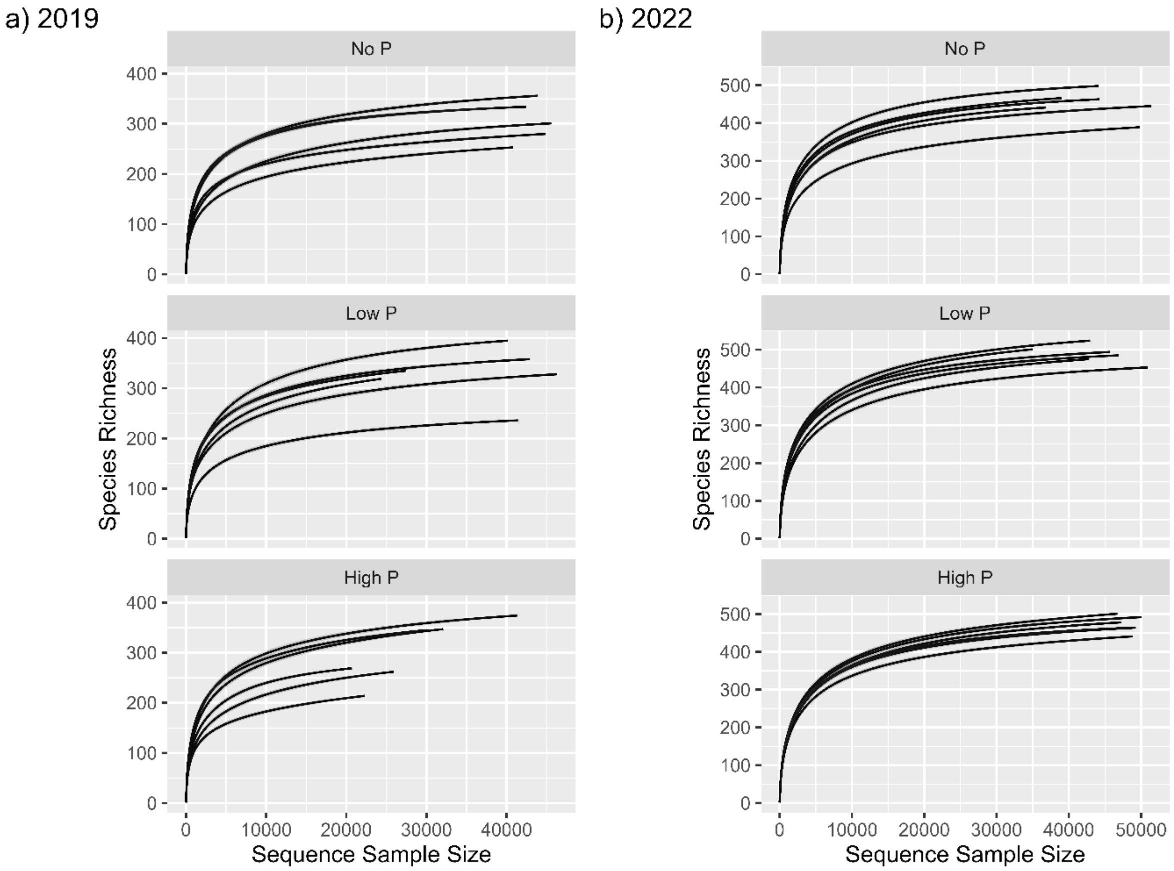


Fig S1. Rarefaction curves for each experimental plot according to long-term P treatment and year.

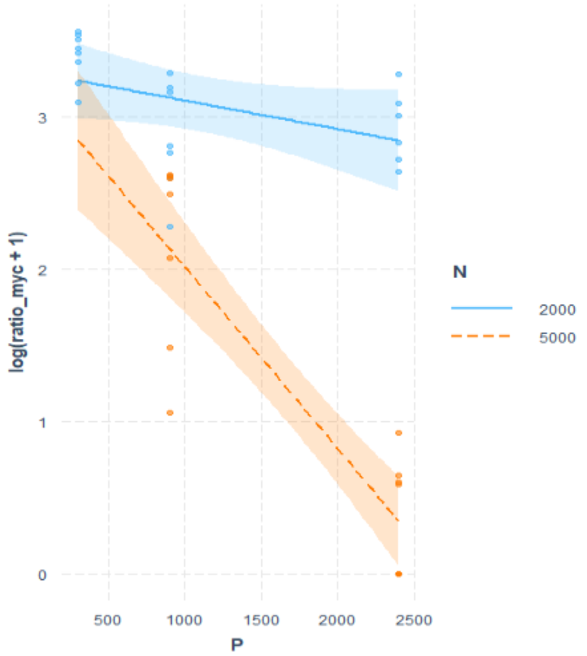


Fig S2. Graph showing the significant interaction between N availability and P availability on mycorrhizal colonisation of wheat (*experiment 3*). In blue N at a concentration of 2000 µM and in orange N at concentration 5000 µM. P range spanning from 500 to 2500 µM. The y axis represents the log transformation of mycorrhizal colonisation. The linear model formula was log(ratio mycorrhiza) = N x P ($r^2_{adj}$= 0.882).

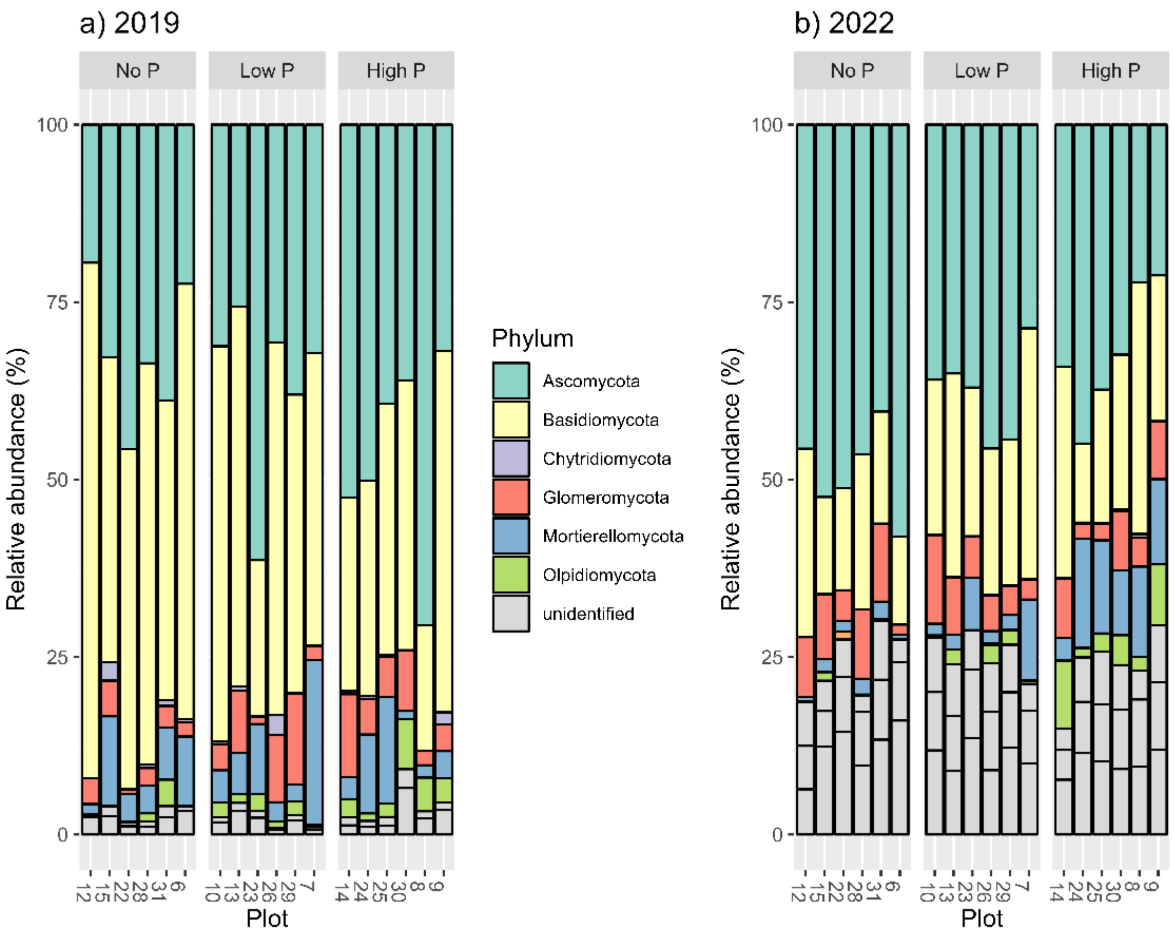


Fig S3. Relative abundance of fungal phylum in wheat roots in plots from a long-term phosphorus fertilisation field trial. Phosphorus fertilisation treatments: No P (no P addition), Low P (replacement P addition, P Olsen equivalent to 2x No P), and High P (P Olsen equivalent to 10 x No P). Wheat was sampled in (a) 2019 and (b) 2022.

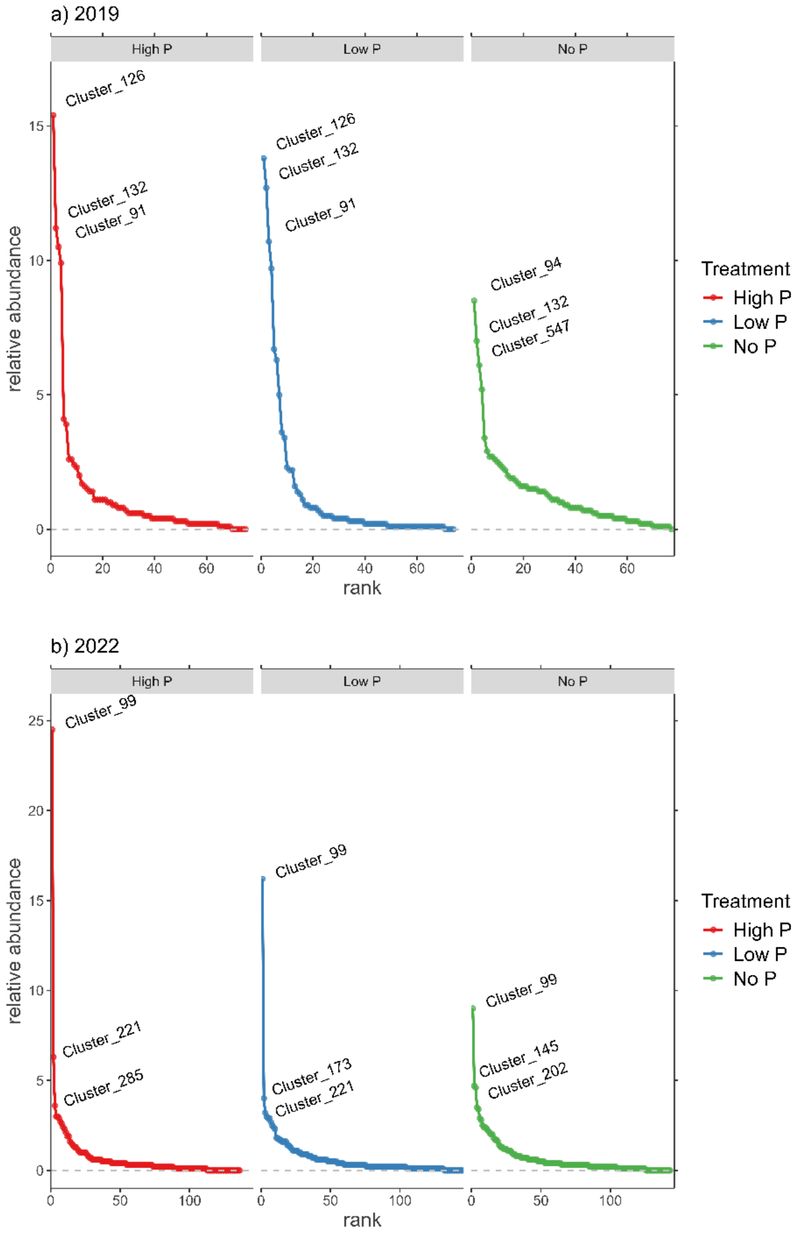


Fig S4. Rank abundance curves of AMF zOTU cluster distribution per year and P treatment. The x-axis represents the ranked zOTUs from high to low. The y-axis shows the average relative abundance for a given zOTU. The three most abundant zOTUs per treatment and year are annotated. Note that cluster definition is independent for 2019 and 2022.

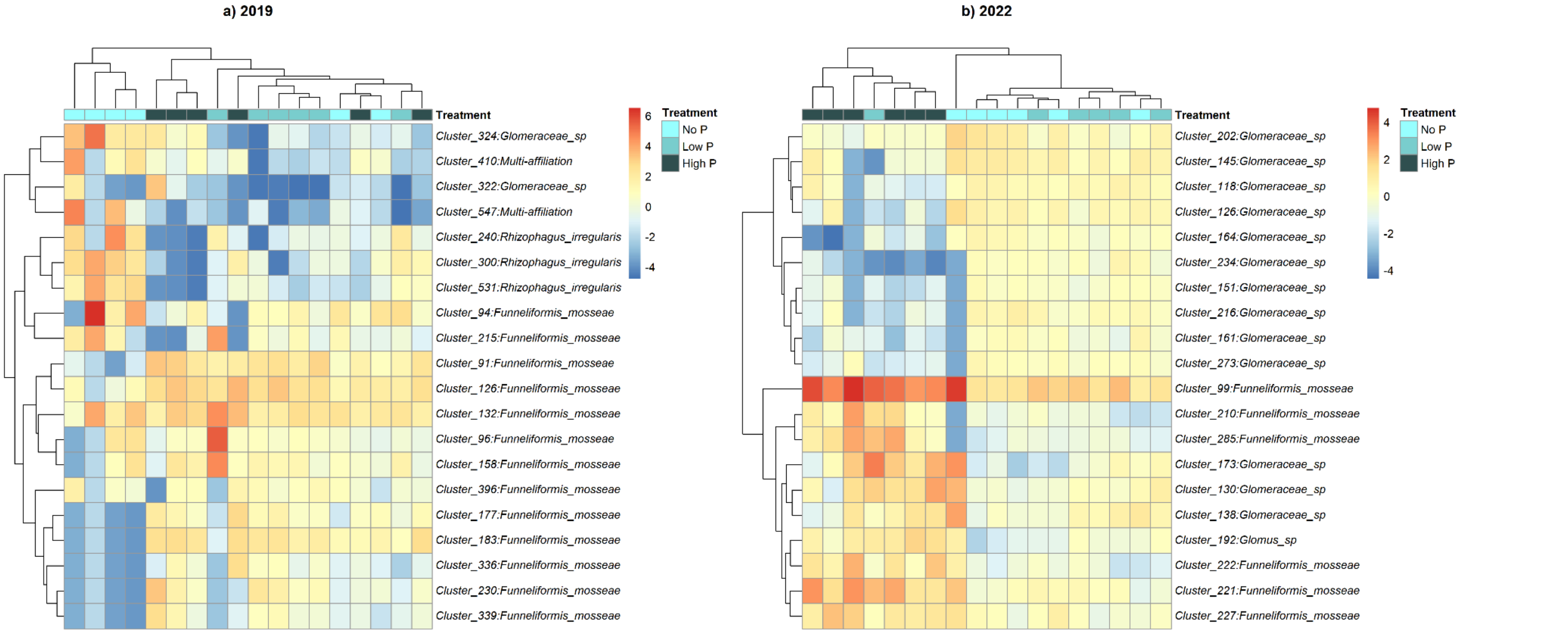


Fig S5. Heatmap showing the root-associated AMF community composition in each plot based on an analysis of the 20 most abundant zOTU clusters. zOTU counts were CLR transformed before analysis. Taxonomic information associated to each zOTU can be found in Supplementary tables S7 & S8. Note that cluster definition is independent for 2019 and 2022.

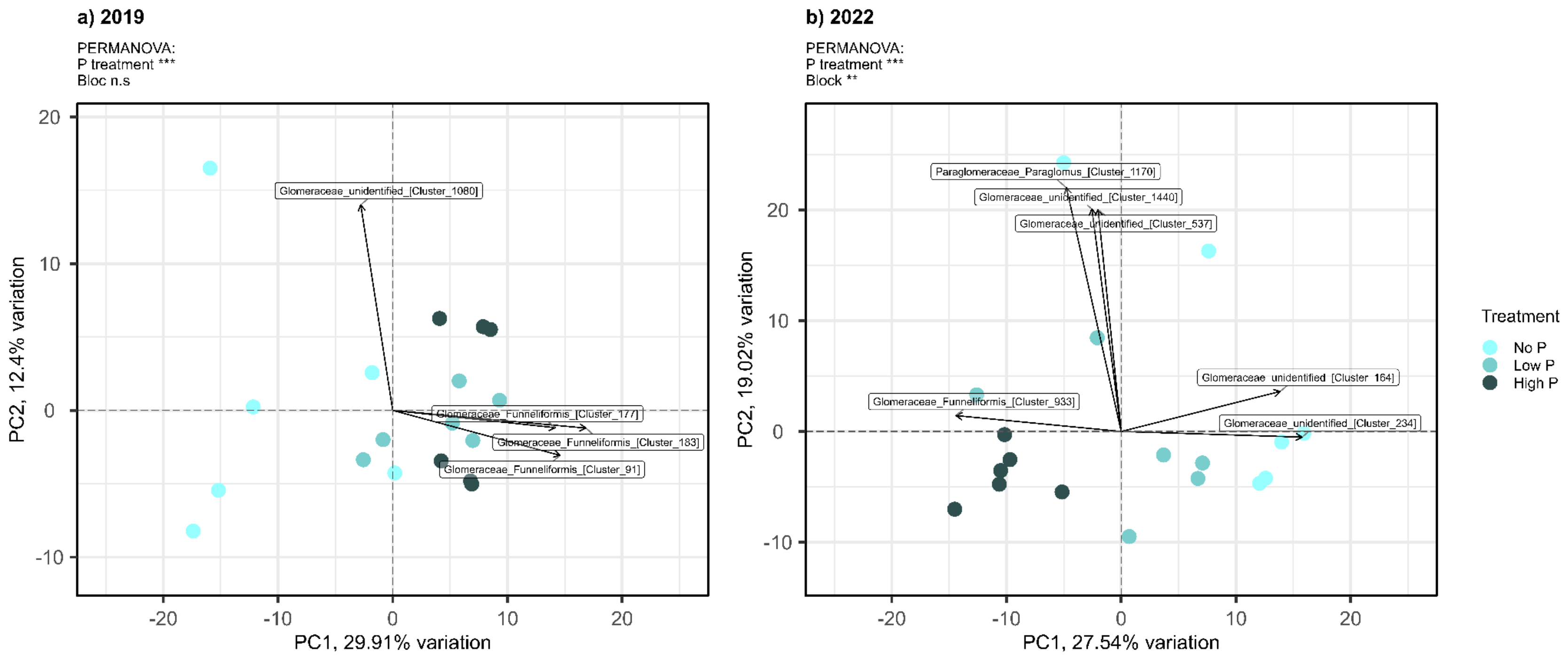


Fig S6. PCAs based on Aitchison distances with CLR transformed AMF zOTU clusters grouped by P treatment. Plots are represented by coloured circles and arrows represent the zOTUs with the greatest loadings in the PCA. Taxonomic information associated to each zOTU (Cluster) can be found in Supplementary tables S7 & S8. Percentages of total variance explained by each principal component (PC1 and PC2) are displayed in the axis titles. Analysis by PERMANOVA detected a significant separation in AMF composition among different P treatments for both years (a) 2029 and (b) 2022. Community differences in centroid location test results: PERMANOVA$_{2019}$: *P treatment* $R^2$= 0.27 $F_{2,13=}$2.84 p<0.01 *block* $R^2$= 0.12 $F_{2,13}$= 1.27 p=0.17; PERMANOVA$_{2022}$: *P treatment* $R^2$= 0.27 $F_{2,13=}$3.28 p<0.001 *block* $R^2$= 0.19 $F_{2,13}$= 2.31 p<0.01. Community differences in multivariate dispersion test results: PERMDISP$_{2019\ F2,15}$= 7.15 p<0.01; PERMDISP$_{2022}$ $F_{2,15}$= 0.23, p=0.83. Pairwise PERMANOVA results among treatments can be found in Table S17.

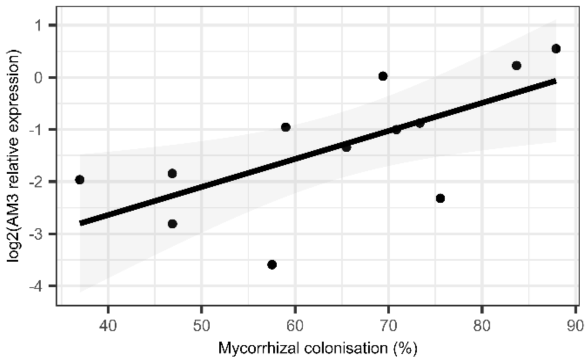


Fig S7: Correlation between the relative expression of the AM3 marker gene and the AMF colonisation level quantified by microscopy using the grid line intersect method. Wheat roots were collected in field during the campaign 2022. $R^2$=0.48.
